# A conserved signaling network monitors delivery of sphingolipids to the plasma membrane in budding yeast

**DOI:** 10.1101/105841

**Authors:** Jesse Clarke, Noah Dephoure, Ira Horecka, Steven Gygi, Douglas Kellogg

## Abstract

In budding yeast, cell cycle progression and ribosome biogenesis are dependent upon plasma membrane growth, which ensures that events of cell growth are coordinated with each other and with the cell cycle. However, the signals that link the cell cycle and ribosome biogenesis to membrane growth are poorly understood. Here, we used proteome-wide mass spectrometry to systematically discover signals associated with membrane growth. The results suggest that membrane trafficking events required for membrane growth generate sphingolipid-dependent signals. A conserved signaling network plays an essential role in signaling by responding to delivery of sphingolipids to the plasma membrane. In addition, sphingolipid-dependent signals control phosphorylation of protein kinase C (Pkc1), which plays an essential role in the pathways that link the cell cycle and ribosome biogenesis to membrane growth. Together, these discoveries provide new clues to how growth-dependent signals control cell growth and the cell cycle.

## Introduction

Cell growth is one of the most fundamental features of life, yet remains poorly understood. Growth is the outcome of multiple processes, including ribosome biogenesis and plasma membrane expansion, that must be precisely coordinated. Numerous questions regarding growth remain unanswered: How are plasma membrane growth and ribosome biogenesis coordinated? How is the rate of growth matched to nutrient availability? How is the amount and location of growth during the cell cycle controlled to maintain a constant size and shape?

In budding yeast, entry into mitosis and ribosome biogenesis are dependent upon plasma membrane growth, which could ensure that growth processes are coordinated with each other and with the cell cycle (Li et al., 2000; Nanduri and Tartakoff, 2001) (Mizuta and Warner, 1994; Anastasia et al., 2012; McCusker and Kellogg, 2012). The linkage was discovered by analyzing the effects of mutants that block membrane trafficking events required for plasma membrane growth. Thus, inactivation of Sec6, which is required for fusion of vesicles with the plasma membrane, causes an arrest of ribosome biogenesis, as well as a pre-mitotic cell cycle arrest. Both pathways signal via a member of the protein kinase C family (Pkc1) that is localized to sites of membrane growth in the daughter bud. Thus, it is possible that key aspects of cell growth are controlled by common signals originating at sites of membrane growth.

The pathway that links mitotic entry to membrane growth has been proposed to control the amount of polar growth that occurs between bud emergence and entry into mitosis, which would influence both cell size and shape (Anastasia et al., 2012). In this model, the vesicles that drive polar membrane growth are thought to deliver signaling molecules that activate Pkc1. As more vesicles fuse with the plasma membrane, a Pkc1-dependent signal would be generated that is proportional to growth, which could be read to determine when sufficient polar growth has occurred. Pkc1 undergoes gradual hyperphosphorylation during polar membrane growth that is dependent upon and proportional to growth, consistent with the idea that it is part of a mechanism that measures polar membrane growth (Anastasia et al., 2012). Growth-dependent signaling suggests a simple and broadly relevant mechanism for control of cell growth and size (McCusker and Kellogg, 2012; Enciso et al., 2014).

The broad outlines of the pathway that links mitotic entry to membrane growth are known. Signaling is dependent upon the Rho1 GTPase, which is delivered to the site of polar growth on vesicles (Abe et al., 2003; Anastasia et al., 2012). Rho1 is activated at the site of growth, where it is thought to bind and activate Pkc1 (Kamada et al., 1996; Abe et al., 2003). Pkc1 binds redundant paralogs called Zds1 and Zds2 that recruit PP2A associated with the Cdc55 regulatory subunit (PP2A^Cdc55^). Pkc1 activates PP2A^Cdc55^, which then activates Mih1, the budding yeast homolog of the Cdc25 phosphatase that drives entry into mitosis by removing Cdk1 inhibitory phosphorylation. Activation of mitotic Cdk1 triggers cessation of polar growth and initiation of isotropic growth that occurs over the entire surface of the bud (Lew and Reed, 1993). Rho1, Pkc1, Zds1/2 and PP2A^Cdc55^ are localized to the site of polar membrane growth and physically interact, providing a direct link between membrane growth and mitotic entry (Yamochi et al., 1994; Kamada et al., 1996; Andrews and Stark, 2000; Rossio and Yoshida, 2011).

A full understanding of how the cell cycle is linked to membrane growth will require a better understanding of the signals generated at sites of membrane growth. A critical question concerns how membrane growth drives phosphorylation and activation of Pkc1. Previous work suggested that the GTP-bound form of Rho1 generated at sites of growth could play a role (Nonaka et al., 1995; Kamada et al., 1996). However, we have thus far been unable to reconstitute hyperphosphorylation of Pkc1 in vitro with purified Rho1-GTP, which suggests that additional signals play a role.

Here, we used proteome-wide mass spectrometry to systematically identify signals associated with membrane growth. To do this, we took advantage of our discovery that Pkc1-dependent signaling rapidly collapses when membrane growth is blocked (Anastasia et al., 2012). Thus, we used proteome-wide mass spectrometry to identify proteins that undergo rapid changes in phosphorylation in response to an arrest of membrane growth.

## Results and Discussion

### Proteome-wide analysis of signals triggered by an arrest of polarized membrane growth

To identify signals triggered by an arrest of polar membrane growth, we released wild type and sec6-4 cells from a G1 arrest and shifted to the restrictive temperature during the polar bud growth phase, which corresponds to the interval of bud growth prior to mitotic entry. Samples for mass spectrometry were taken 5 minutes after inactivation of sec6-4. Western blotting was used to confirm that the wild type and sec6-4 cells were at the same point in the cell cycle, and that inactivation of sec6-4 caused rapid loss of Pkc1 phosphorylation, as previously observed (Anastasia et al., 2012). Proteolytic peptides from each strain were phospho-enriched, covalently modified by reductive dimethylation to generate light (wild type) and heavy (sec6-4) stable isotope labeled pools, and then combined and analyzed by LC-MS/MS (Villén and Gygi, 2008; Kettenbach and Gerber, 2011). The heavy to light ratios of phosphorylated peptides in sec6-4 cells versus wild type cells were log_2_ transformed. Thus, negative values indicate decreased phosphorylation in sec6-4 cells, while positive values indicate increased phosphorylation. Three biological replicates were analyzed, which allowed calculation of average log_2_ ratios and standard deviations for most peptides.

The complete data set contains >75,000 spectral matches filtered to <0.1% peptide false discovery rate and 1% protein false discovery rate using the target-decoy approach (see methods). Table S1 presents a summary of all identified phosphorylation sites with quantitative data. Table S2 provides detailed information for each of the detected phosphopeptides. A total of 9375 sites were identified on 1831 proteins. Of these, 6375 sites on 1694 proteins could be quantified. The overlap of quantified sites between the three biological replicates is shown in Figure Supplement 1.

We focused on sites that were quantified in at least two of the three biological replicates, which comprised 4485 sites on 1520 proteins. Within this set, we focused on sites with a median log_2_ ratio greater than 1.2 in either direction. At this threshold, 69 sites on 58 proteins appeared to undergo rapid dephosphorylation when *sec6-4* was inactivated (Table S3). 63 sites on 57 proteins appeared to undergo an increase in phosphorylation (Table S4). Both Pkc1 and Zds1 were detected as undergoing dephosphorylation in sec6-4 cells (Table S3). Since these proteins were previously shown to undergo dephosphorylation after inactivation of Sec6, their presence in the data provides evidence that the approach can identify relevant proteins (Anastasia et al., 2012). Inactivation of Sec6 may cause effects that are independent of its role in membrane growth. For simplicity, however, we refer to the effects of inactivating Sec6 as a block to membrane growth.

### Blocking membrane traffic disrupts a signaling network that controls cell growth

We searched the mass spectrometry data for proteins that were previously linked to cell growth and discovered that multiple components of a target of rapamycin complex 2 (TORC2) signaling network undergo changes in phosphorylation in response to an arrest of membrane growth (Tables S3 and S4). The TORC2 network controls a pair of partially redundant kinase paralogs called Ypk1 and Ypk2, which are homologs of vertebrate SGK (Casamayor et al., 1999). Extensive work has shown that TORC2 and Ypk1/2, as well as their surrounding signaling network, play roles in controlling cell growth, lipid synthesis and cell cycle progression. In addition, multiple previous studies have suggested that TORC2 and Ypk1/2 control Rho1/Pkc1 signaling (Helliwell et al., 1998; Roelants et al., 2002; Schmelzle et al., 2002; Kamada et al., 2005; Niles and Powers, 2014; Hatakeyama et al., 2017).

Figure 1A provides an overview of the TORC2-Ypk1/2 network, with proteins identified by mass spectrometry highlighted in red. Ypk1/2 are phosphorylated by TORC2 and phosphoinositide-dependent kinase 1 (PDK1), which play conserved roles in controlling cell growth (Roelants et al., 2002; Kamada et al., 2005). In budding yeast there are two redundant homologs of PDK1 called Pkh1 and Pkh2. TORC2 and Pkh1/2 phosphorylate distinct sites on Ypk1/2 that are thought to increase activity.

**Figure 1.**
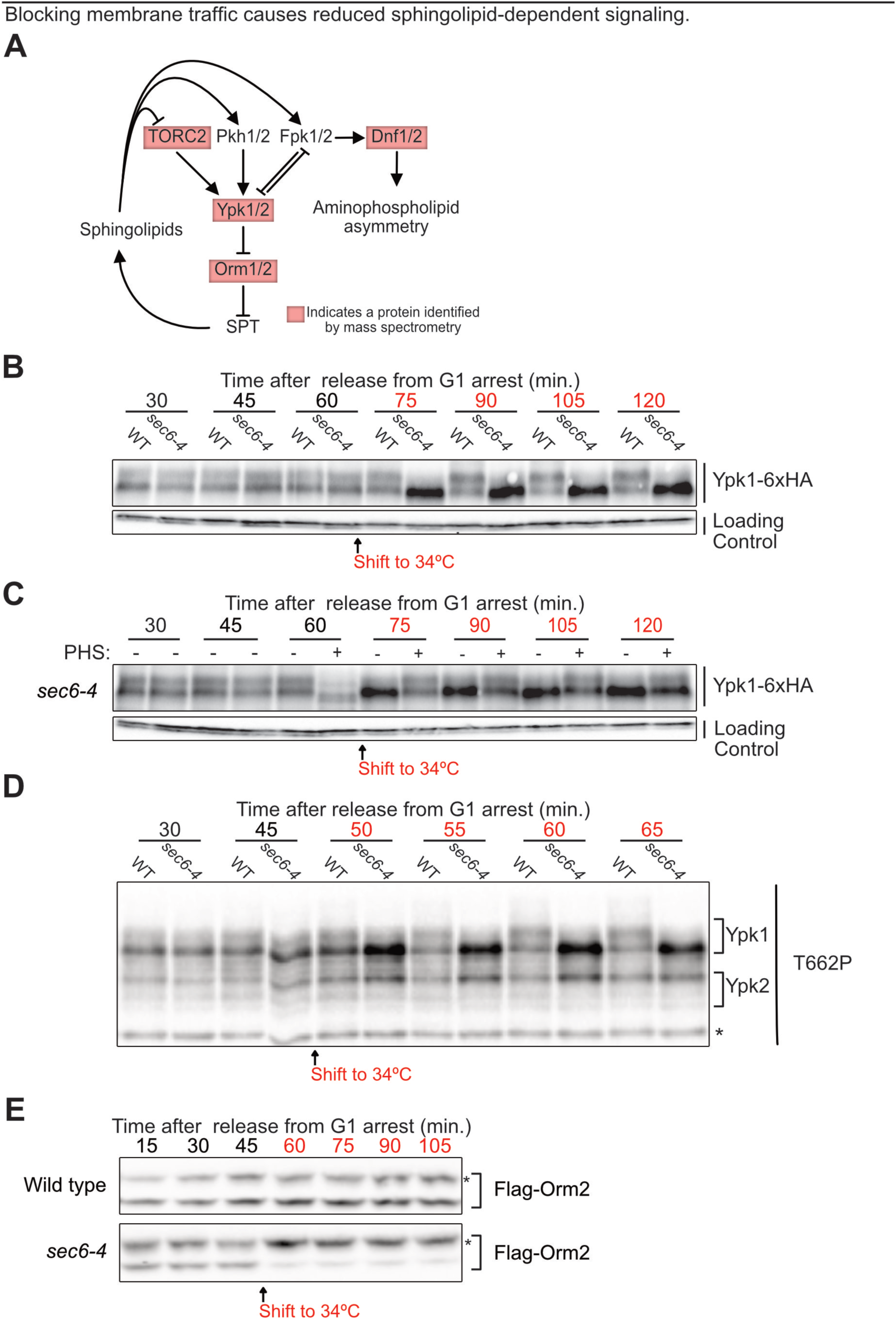
(**A**) Proteins in the TORC2-Ypk1/2 signaling network that showed a large change in phosphorylation upon growth arrest are highlighted in red. (**B**)YPK1-6xHA cells with or without the sec6-4 allele were released from G1 arrest at 22°C and shifted to 34°C after 60 minutes (when cells reached ∼50% budding). Ypk1-6xHA phosphorylation was assayed by western blot. The same blot was probed for Nap1 to provide a loading control. (**C**) YPK1-6xHA sec6-4 cells were released from G1 arrest at 22°C and shifted to 34°C after 60 minutes (when cells reached ∼50% budding). 10 *µ*M phytosphingosine was added to one half of the culture (+) immediately before the 60 minute time point. Ypk1 phosphorylation was assayed by western blot. The slight increase in electrophoretic mobility of Ypk1-6XHA immediately after addition of phytosphingosine at 60 minutes was also observed in wild type control cells, where it disappeared within 5 minutes (not show). The reason for the transient change in electrophoretic mobility is unknown. The same blot was probed for Nap1 to provide a loading control. (**D**) Cells with or without the sec6-4 allele were released from G1 arrest at 22°C and shifted to 34°C after 45 minutes (when approximately 10% of cells had buds). TORC2-dependent phosphorylation of Ypk1-T662P was assayed by western blot with a phospho-specific polyclonal antibody that recognizes TORC2 sites on Ypk1 and Ypk2. A non-specific background band that serves as a loading control is marked with an asterisk. (**E**) Flag-ORM2 cells with or without the sec6-4 allele were released from G1 arrest at 22°C and shifted to 34°C after the 45 minute time point (when approximately 10% of cells had buds). Flag-Orm2 phosphorylationwas assayed by running samples on a Phos-Tag gel followed by western blot. Hyperphosphorylated forms of Orm2 are marked with an asterisk.

One function of the network is to modulate synthesis of sphingolipids, a diverse family of lipids that includes components of the plasma membrane, as well as lipids that play roles in signaling (Aronova et al., 2008; Dickson, 2008; Roelants et al., 2011; Muir et al., 2014). Sphingolipid synthesis is initiated at the endoplasmic reticulum by serine palmitoyltransferase (SPT), which synthesizes precursors for production of phytosphingosine and ceramides, which are used to produce complex sphingolipids in the Golgi. Complex sphingolipids are transported to the plasma membrane and together constitute approximately 10% of membrane lipids (Ejsing et al., 2009; Klose et al., 2012). Ypk1/2 promote sphingolipid synthesis by phosphorylating and inhibiting Orm1 and Orm2, redundant paralogs that bind and inhibit serine palmitoyltransferase (Breslow et al., 2010; Roelants et al., 2011).

Ypk1/2 are also phosphorylated by a pair of redundant kinase paralogs called Fpk1 and Fpk2 that localize to the site of polar growth (Nakano et al., 2008; Roelants et al., 2010). Genetic data suggest that complex sphingolipids are required for Fpk1/2 activity, and that Fpk1/2 and Ypk1/2 inhibit each other (Roelants et al., 2010). Fpk1/2 also phosphorylate a redundant pair of lipid flippases called Dnf1 and Dnf2, which are located at the site of polar growth and are thought to flip phospholipids from the outer to inner leaflet of the plasma membrane (Hua et al., 2002; Pomorski et al., 2003; Nakano et al., 2008). Loss of FPK1/2 or DNF1/2 causes mild elongation of the daughter bud, which suggests that they play a role in controlling polar growth (Pomorski et al., 2003; Nakano et al., 2008).

### Blocking membrane traffic causes reduced sphingolipid-dependent signaling

Multiple proteins in the TORC2-Ypk1/2 network underwent significant changes in phosphorylation in response to inactivation of Sec6, including Ypk1, Orm2, Dnf2 and Avo1, a component of the TORC2 complex (Tables S3 and S4). All of these proteins regulate and/or respond to sphingolipids. To further investigate changes in the network, we first analyzed phosphorylation of Ypk1. Ypk1 undergoes Fpk1/2-dependent phosphorylation that causes a decrease in electrophoretic mobility that can be detected by western blotting (Roelants et al., 2011). Inactivation of Sec6 caused loss of Fpk1/2-dependent phosphorylation of Ypk1, which suggests that blocking membrane traffic causes inactivation of Fpk1/2 (Figure 1B). Dnf2, a direct target of Fpk1/2, also showed a loss of phosphorylation in the mass spectrometry data, which provided further evidence for a loss of Fpk1/2 activity (Table S3). Since Fpk1/2 activity is dependent upon complex sphingolipids, these observations suggest that inactivation of Sec6 causes a decrease in sphingolipid-dependent signaling. Addition of exogenous phytosphingosine rescued the loss of Ypk1 phosphorylation caused by arresting growth, consistent with the idea that inactivation of Sec6 causes loss of sphingolipid-dependent signals (Figure 1C). The rescue faded over longer times, which suggests that the relevant signaling lipids are short-lived or transported to a location where they can not influence Fpk1/2 activity.

Previous studies have suggested that loss of sphingolipids causes increased TORC2 activity (Roelants et al., 2011; Berchtold et al., 2012). Moreover, the mass spectrometry analysis found that Avo1, a component of the TORC2 complex, undergoes increased phosphorylation in response to an arrest of membrane growth (Table S3). Since Avo1 is a target of TORC2, this observation suggests that TORC2 activity increases (Wullschleger et al., 2005). To test this, we assayed TORC2-dependent phosphorylation of Ypk1/2 using a phosphospecific antibody. Inactivation of Sec6 caused increased TORC2-dependent phosphorylation of Ypk1 (Figure 1D). We again observed that inactivation of Sec6 caused loss of the Fpk1/2-dependent mobility shift of Ypk1.

The site on Orm2 that was detected (threonine 18) was previously identified as a likely Ypk1/2 target (Breslow et al., 2010; Roelants et al., 2010; 2011). The mass spectrometry data suggested that Orm2 is dephosphorylated when Sec6 is inactivated (Table S4). To investigate further, we used Phos-tag gels and western blotting to assay Orm2 phosphorylation after inactivation of Sec6. In contrast to expectations from the mass spectrometry data, Orm2 was rapidly hyperphosphorylated when Sec6 was inactivated, which suggests that Ypk1/2 become more active (Figure 1E). The disagreement between the mass spectrometry and western blotting data may be a product of the complexity of phosphopeptide analysis. Phosphosites on Orm1/2 are located close to each other, such that multiple sites are found on the same proteolytic peptide. If a singly phosphorylated peptide is converted to a multiply phosphorylated peptide upon inactivation of Sec6, the analysis will detect a decrease in the abundance of the singly phosphorylated peptide, giving the appearance of a dephosphorylation event.

The Orm1/2 proteins are also phosphorylated by the kinase Npr1. To test whether hyperphosphorylation of Orm2 was due to Ypk1/2, we utilized sec6-4 cells that are dependent upon an analog-sensitive allele of Ypk1 (sec6-4 ypk1-as *ypk2*Δ), which allows rapid inactivation of Ypk1/2 with the inhibitor 3-MOB-PP1 (Sun et al., 2012). Inactivation of Ypk1/2 blocked hyperphosphorylation of Orm2 when membrane growth was blocked (Figure Supplement 2A). In addition, a mutant version of Orm2 that lacks key sites phosphorylated by Ypk1/2 also failed to undergo hyperphosphorylation (Figure Supplement 2B). Thus, hyperphosphorylation of Orm2 in response to inactivation of Sec6 is dependent upon Ypk1/2.

Since the Orm1/2 proteins are embedded in the membrane of the endoplasmic reticulum (Han et al., 2010), the discovery that they undergo rapid hyperphosphorylation in response to an arrest of plasma membrane growth suggests that there is rapid communication between the plasma membrane and the endoplasmic reticulum. Approximately 50% of the endoplasmic reticulum is closely associated with the plasma membrane, which would facilitate rapid communication (Pichler et al., 2001).

Previous work found that inhibition of the first step of sphingolipid synthesis causes inactivation of Fpk1/2, activation of both TORC2 and Ypk1/2, and hyperphosphorylation of Orm1/2 (Breslow et al., 2010; Roelants et al., 2011; Berchtold et al., 2012; Sun et al., 2012). Since hyperphosphorylation of Orm1/2 relieves inhibition of sphingolipid synthesis, the TORC2-Ypk1/2 network was proposed to monitor sphingolipid synthesis to maintain sphingolipid homeostasis (Breslow et al., 2010). However, the mechanism by which sphingolipid synthesis influences signaling in the network has remained unknown. Here, we found that blocking traffic to the plasma membrane causes effects identical to those caused by inhibition of sphingolipid synthesis. In addition, although inactivation of Sec6 caused hyperphosphorylation of Orm2, which should stimulate sphingolipid synthesis at the endoplasmic reticulum, there was no evidence that increased sphingolipid synthesis was perceived by the network: TORC2-dependent phosphorylation of Ypk1/2 persisted, Fpk1/2 remained inactive, and Orm2 phosphorylation persisted. Together, these observations suggest that the signaling network monitors delivery of sphingolipids to the plasma membrane by vesicles, rather than synthesis of sphingolipids at the endoplasmic reticulum.

A number of previous studies support the idea that sphingolipids influence signaling events at the plasma membrane. For example, TORC2 is localized to plasma membrane foci, so it is positioned to respond to signals at the plasma membrane (Berchtold and Walther, 2009). Furthermore, inhibition of sphingolipid synthesis triggers release of the Slm1 and Slm2 proteins from plasma membrane compartments called eisosomes (Berchtold et al., 2012). Slm1/2 promote TORC2-dependent activation of Ypk1/2 (Niles et al., 2012). Thus, the localization of TORC2 and Slm1/2, and their behavior in response to inhibition of sphingolipid synthesis, are consistent with the idea that they respond to sphingolipid-dependent signals at the plasma membrane. However, since the plasma membrane is closely associated with the endoplasmic reticulum, the localization and behavior of TORC2 and Slm1/2 alone would not rule out the possibility that they directly monitor synthesis of sphingolipids at the endoplasmic reticulum.

A model that could explain the data is that post-Golgi vesicles required for polar growth deliver sphingolipids to the plasma membrane. When vesicle delivery is blocked, sphingolipids at the plasma membrane rapidly decline, which could occur by modification or sequestration into lipid domains. Loss of sphingolipids, in turn, would lead to inactivation of Fpk1/2 and a consequent loss of Ypk1/2 and Dnf1/2 phosphorylation. A decrease in sphingolipids would also cause increased activity of TORC2, which stimulates Ypk1/2 to hyperphosphorylate Orm1/2. The Ypk1/2 signaling network has also been proposed to respond to membrane stress (Berchtold et al., 2012). Thus, an alternative model is that blocking delivery of vesicles to the plasma membrane triggers membrane stress that activates the pathway.

### Inactivation of Ypk1/2 causes a failure in Pkc1 phosphorylation that is rescued by exogenous sphingolipids

We next used ypk1-as *ypk2*Δ cells to test whether Ypk1/2 are required for normal hyperphosphorylation of Pkc1 during polar membrane growth. Since hyperphosphorylation of Pkc1 is dependent upon and proportional to polar membrane growth, it provides a readout of signaling events associated with membrane growth. Wild type control cells and ypk1-as *ypk2*Δ cells were released from G1 arrest and 3-MOB-PP1 was added to both cultures before cells had initiated bud emergence. Addition of inhibitor to ypk1-as *ypk2*Δ cells caused a failure in Pkc1 hyperphosphorylation (Figure 2A), as well as a failure in mitotic entry that was detected as a failure to produce the mitotic cyclin Clb2 (Figure 2B). It also caused delayed bud emergence (Figure 2C). In some experimental replicates, Pkc1 phosphorylation was strongly reduced, but not completely eliminated (see, for example, Figure 3A). The fact that bud emergence still occurred when Ypk1/2 were inhibited indicated that the failure in Pkc1 phosphorylation was not due simply to a failure in bud growth. To determine whether the delay in bud emergence was due to decreased Pkc1 activity, we expressed a constitutively active version of PKC1 (PKC1*) in ypk1-as *ypk2*Δ cells (Watanabe et al., 1994). We used basal expression from the uninduced CUP1 promoter to achieve low level expression of PKC1* (Thai et al., 2017). Expression of PKC1* rescued the delay in bud emergence (Figure 2C). Together, these observations suggest that inhibition of Ypk1/2 causes decreased phosphorylation of Pkc1, as well as decreased Pkc1 activity.

**Figure 2.**
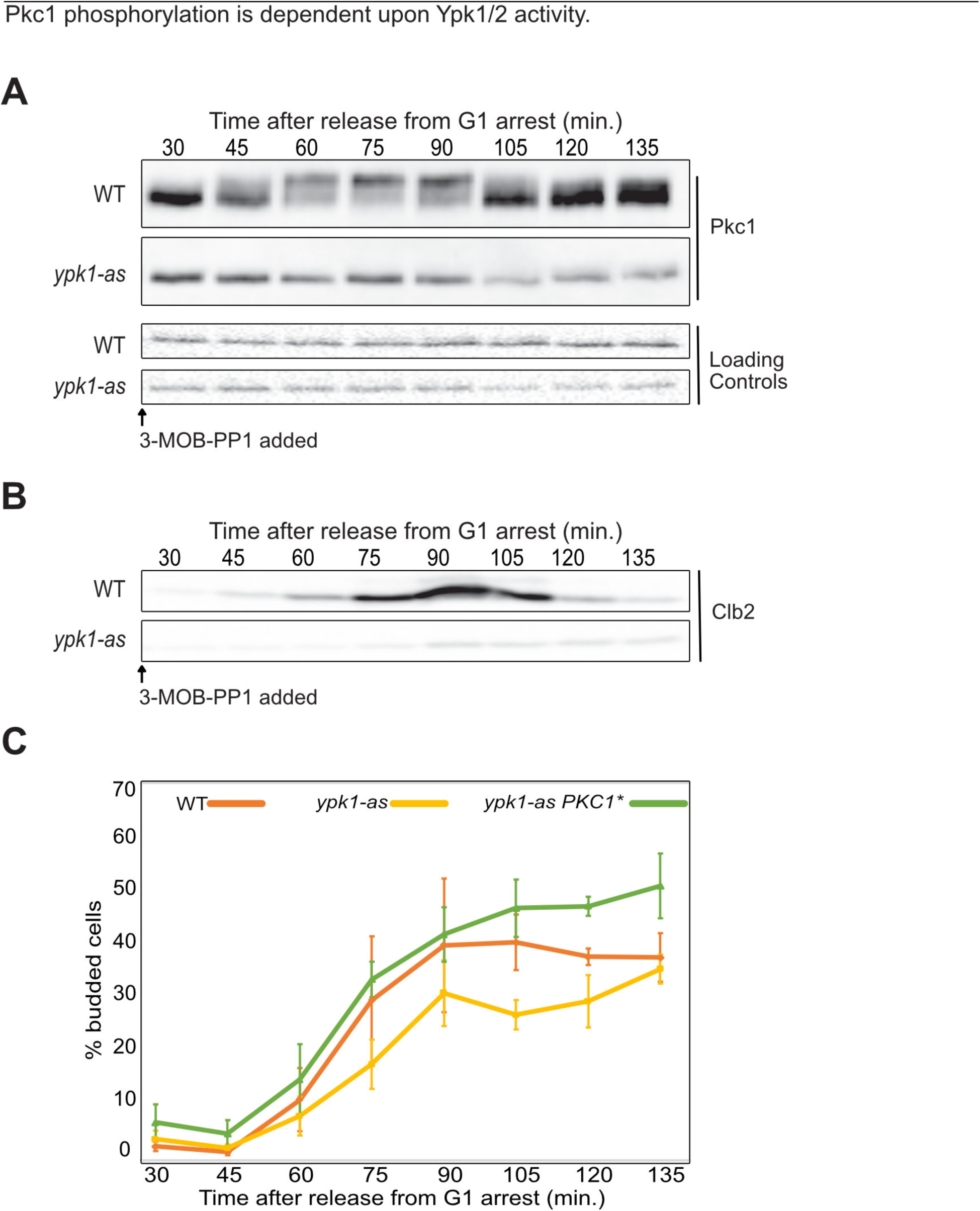
(**A**) The indicated cells were released from a G1 arrest at 25°C and 50 *µ*M 3-MOB-PP1 was added to each strain at 10 min after release. Pkc1 phosphorylation was assayed by western blot. A background band from the same blot was used as a loading control. (**B**) A western blot showing levels of the mitotic cyclin Clb2 from the same samples as panel A. (**C**) Cells were released from a G1 arrest at 25°C and 50 *µ*M 3-MOB-PP1 was added at 10 min after release. The percentage of budded cells was determined by counting the number of budded cells in a total of at least 200 cells for each time point. The percentage of budded cells was averaged for three biological replicates. Error bars represent the standard error of the mean for three biological replicates.

**Figure 3.**
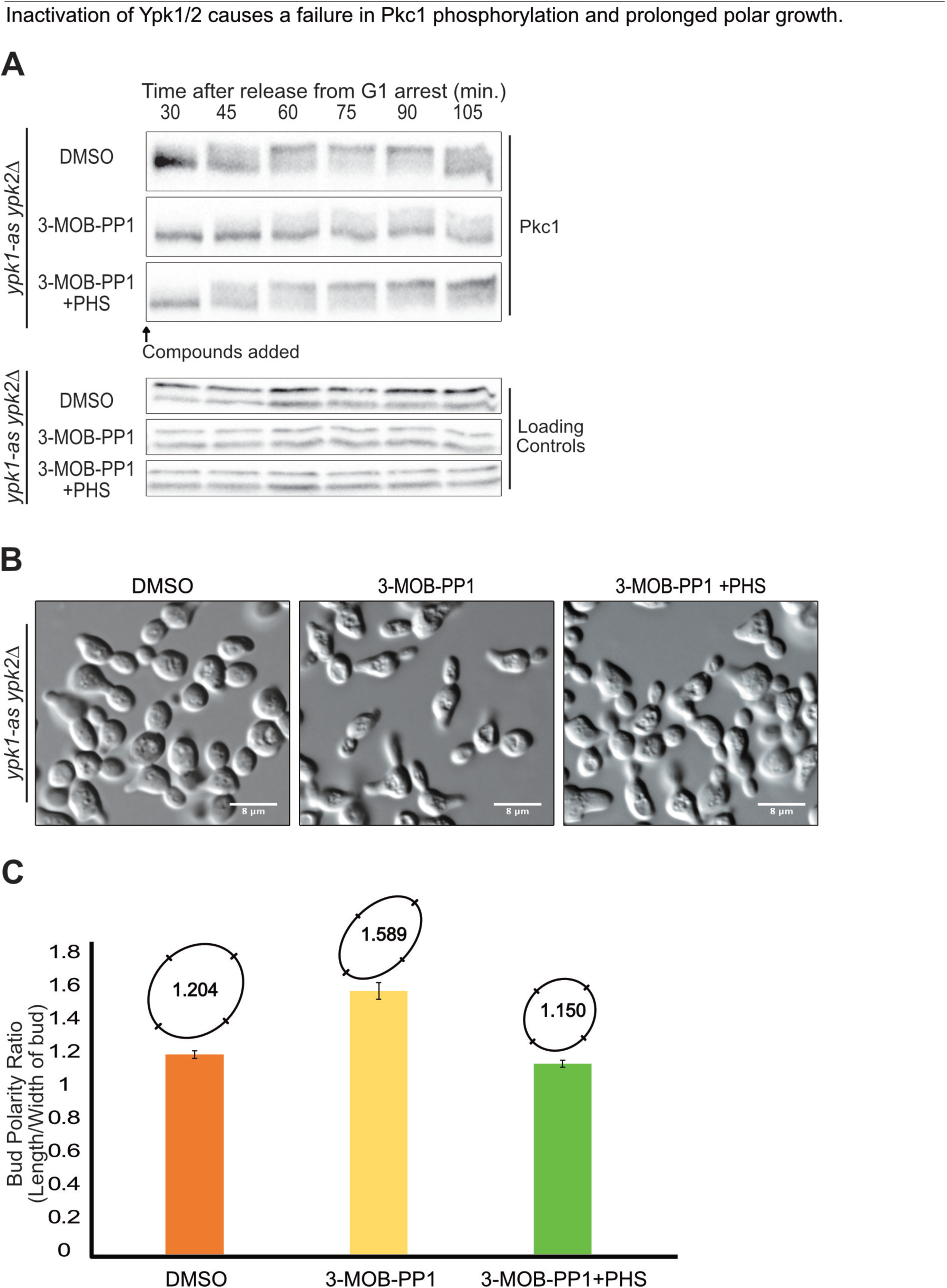
(**A**) ypk1-as *ypk2*Δ cells were released from a G1 arrest at 25°C and the indicated compounds were added at 10 min after release. Pkc1 phosphorylation was assayed by western blot. A background band from the same blot was used as a loading control. **(B**) Fixed cells from the 120 minute time point of panel A were imaged by DIC microscopy. White bars are 8 *µ*m. (**C**) Bud polarity of the imaged cells in panel B was determined by calculating the ratio of the bud length to width of buds for 50 cells and then plotting the average ratio. Error bars represent the standard error of the mean. The average polarity ratios were also converted to ovals whose relative shape can be directly compared.

Inhibition of Ypk1/2 should lead to decreased synthesis of sphingolipids due to hyperactivity of Orm1/2 (Breslow et al., 2010; Roelants et al., 2011; Sun et al., 2012). We therefore added exogenous phytosphingosine to test whether failure to phosphorylate Pkc1 when Ypk1/2 are inhibited is a consequence of decreased sphingolipid production. Addition of phytosphingosine largely restored hyperphosphorylation of Pkc1 when Ypk1/2 were inhibited before bud emergence (Figure 3A, compare the fraction of Pkc1 that reaches the fully hyperphosphorylated form).

The buds that emerged during Ypk1/2 inhibition were elongated (Figure 3B and Figure Supplement 2C). Cell elongation was quantified by measuring the average ratio of bud length to width (Figure 3C). Addition of phytosphingosine largely eliminated the elongated bud phenotype (Figure 3B, C). Previous studies showed that bud elongation can be a consequence of a decrease in the activities of Pkc1 or mitotic Cdk1 (Lew and Reed, 1993; Anastasia et al., 2012).

Although inhibition of Ypk1/2 in G1 phase blocked Pkc1 phosphorylation, inhibition of Ypk1/2 later in the cell cycle, during polar bud growth, did not cause loss of Pkc1 phosphorylation (Figure Supplement 3A). Thus, Pkc1 phosphorylation is independent of Ypk1/2 activity during polar growth. This observation suggests that Ypk1/2 are unlikely to control Pkc1 phosphorylation directly, and that rapid dephosphorylation of Pkc1 caused by inactivation of Sec6 during polar bud growth is unlikely to be due to stimulation of the TORC2-Ypk1/2 network. Rather, Ypk1/2 could promote Pkc1 phosphorylation by stimulating production of sphingolipids earlier in the cell cycle.

Since the relationship between Pkc1 phosphorylation and Pkc1 kinase activity is poorly understood, these data do not establish whether sphingolipids promote Pkc1 activity. However, previous studies found that defects caused by inhibition of serine palmitoyltransferase can be suppressed by increased activity of Pkc1, which suggests that inhibition of sphingolipid synthesis leads to a failure to fully activate Pkc1 (Friant et al., 2000; Olson et al., 2015). There is also evidence that sphingolipids promote Pkc1 activity via activation of the Rho1 GTPase, although the underlying mechanisms remain largely unknown (Niles and Powers, 2014; Olson et al., 2015; Hatakeyama et al., 2017). Together, the data suggest that sphingolipids promote Pkc1 activation, potentially by promoting Pkc1 phosphorylation.

Sphingolipids are also thought to regulate Pkh1/2, which directly phosphorylate Pkc1 on a site that stimulates its activity (Inagaki et al., 1999; Sun et al., 2000; Friant et al., 2001; Roelants et al., 2004; Liu et al., 2005). Since Pkh1/2 phosphorylate Pkc1 on a single site, they are unlikely to be directly responsible for the gradual multi-site hyperphosphorylation of Pkc1 that occurs during polar membrane growth. However, Pkh1/2 could activate Pkc1 to undergo autophosphorylation. To test for a contribution of Pkh1/2, we utilized an analog-sensitive allele of PKH1 in a *pkh2*Δ background (pkh1-as *pkh2*Δ) (Sun et al., 2012). Inactivation of Pkh1/2 during polar growth caused a rapid and complete loss of Pkh1/2-dependent phosphorylation of Ypk1, but did not cause a loss of Pkc1 phosphorylation (Figure 4A). These results indicate that the rapid dephosphorylation of Pkc1 observed in response to an arrest of polar membrane growth is not likely to be caused by loss of Pkh1/2 activity. We also found that hyperphosphorylation of Pkc1 occurred normally in *fpk1*Δ *fpk2*Δ cells, which rules out a model in which loss of Pkc1 phosphorylation is due to loss of Fpk1/2 activity (Figure Supplement 3B).

**Figure 4.**
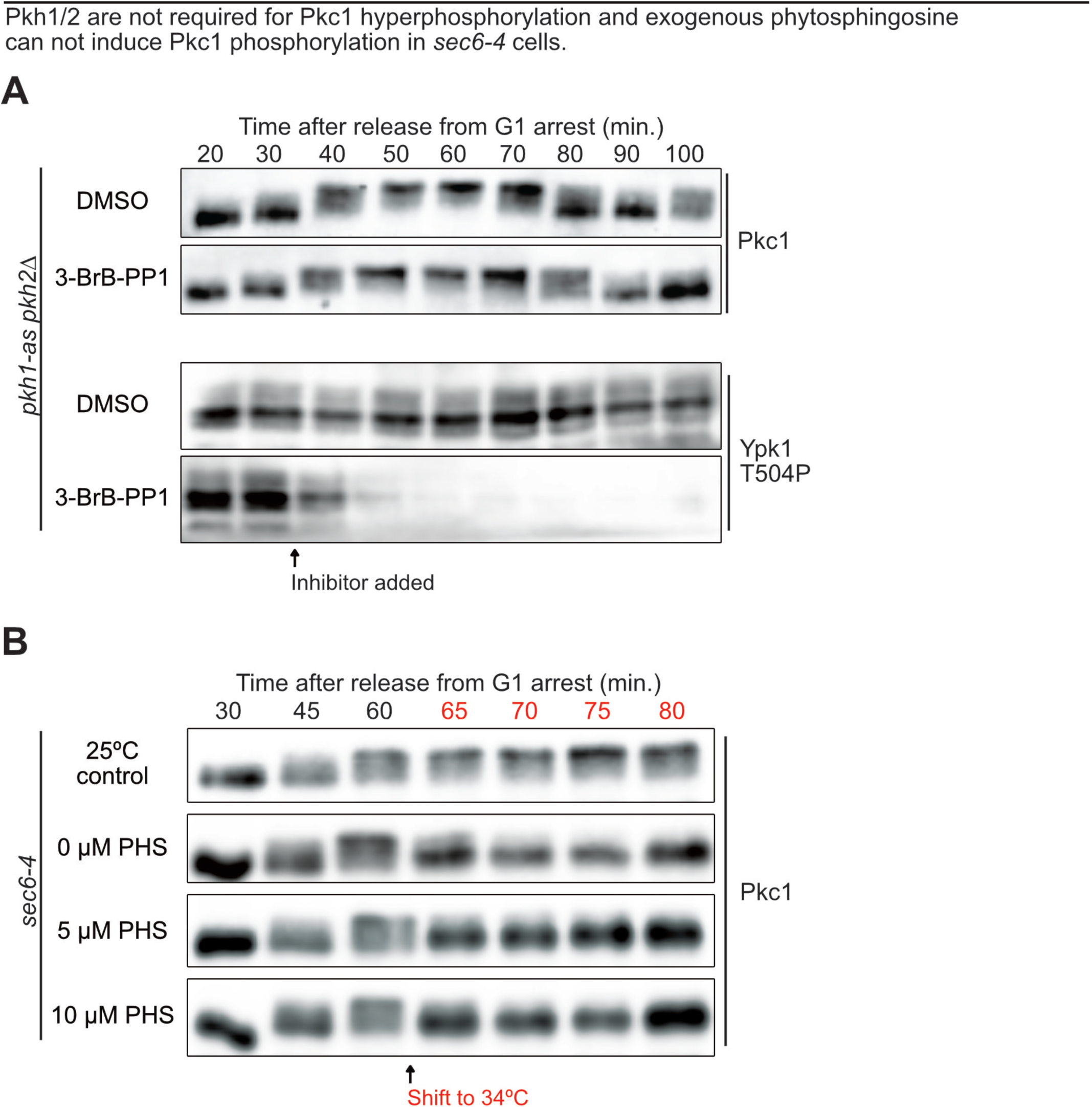
(**A**) pkh1-as *pkh2*Δ were released from a G1 arrest and DMSO or 15 *µ*M 3-BrB-PP1 were added at 30 min after release. Pkc1 phosphorylation was assayed by western blot. Pkh1/2-dependent phosphorylation of Ypk1-T504 was assayed by western blot using a phosphospecific antibody. Blots are from the same samples so timing can be directly compared. (**B**) sec6-4 cells were released from a G1 arrest at 25°C. At 60 minutes the culture was split into 4 aliquots and the indicated amounts of phytosphingosine were added to cultures shifted to 34°C. Pkc1 phosphorylation was assayed by western blot.

Addition of exogenous phytosphingosine did not rescue loss of Pkc1 phosphorylation caused by inactivation of Sec6 (Figure 4B). This observation is consistent with a model in which vesicle-dependent delivery of sphingolipids to the plasma membrane is required for sphingolipids to influence Pkc1 phosphorylation.

### Inhibition of sphingolipid production causes reduced phosphorylation of Pkc1

To further define the contribution of sphingolipids, we utilized a temperature sensitive allele of LCB1 (lcb1-100), which encodes an essential subunit of serine palmitoyltransferase (Meier et al., 2006). **I**nactivation of Lcb1 caused a decrease in Pkc1 hyperphosphorylation during polar growth, which could be seen as a failure to produce the most hyperphosphorylated forms of Pkc1 (Figure 5). Analysis of mitotic cyclin levels revealed that inactivation of Lcb1 also caused delayed mitotic entry and a prolonged mitosis (Figure 5). Cells budded when Lcb1 was inactivated so the delay was not due to a failure in bud growth. Addition of exogenous phytosphingosine partially rescued Pkc1 hyperphosphorylation and fully rescued the mitotic delay, which suggests that the effects of lcb1-100 were due at least partly to a failure to produce sphingolipids (Figure 5).

**Figure 5.**
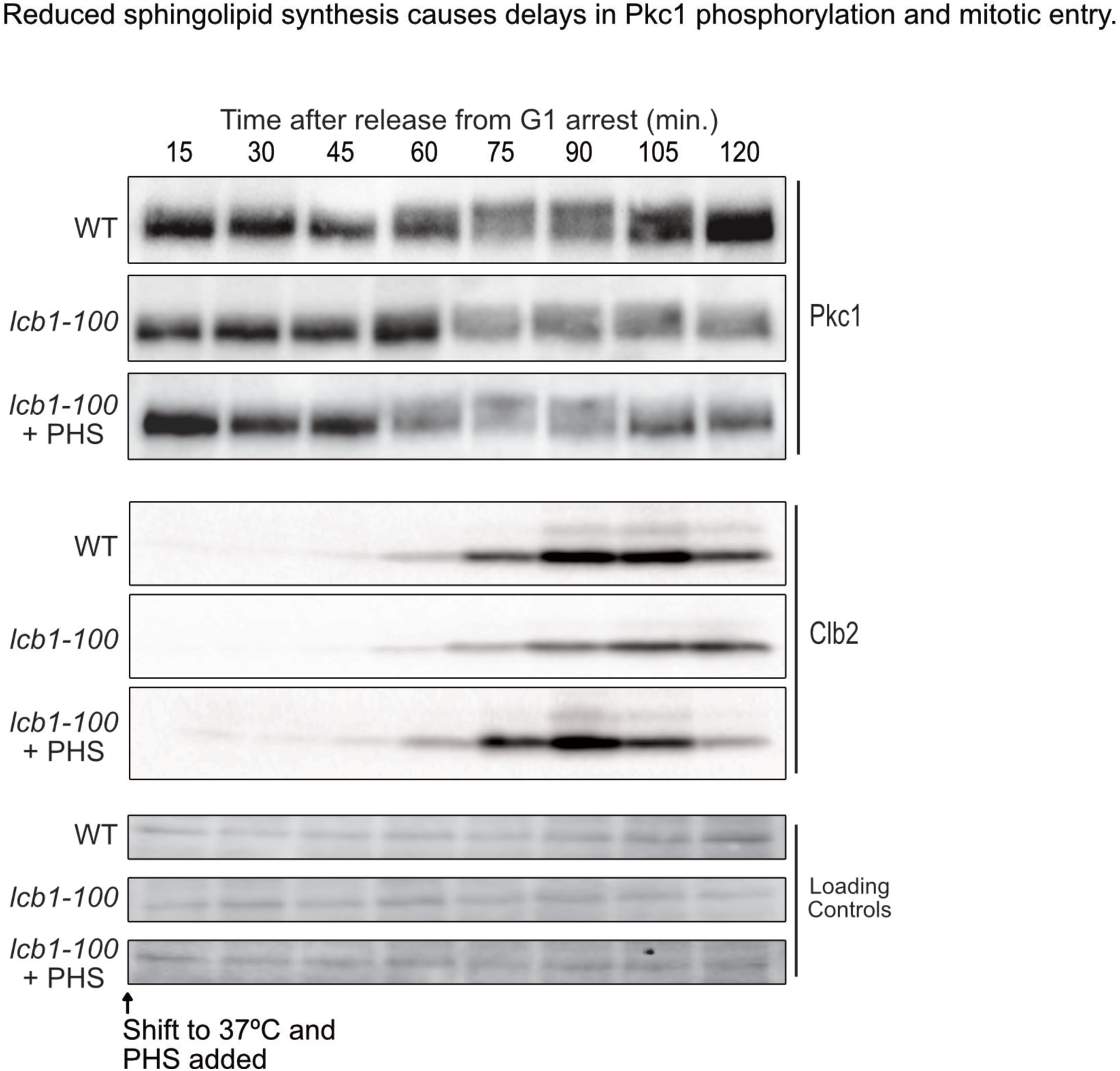
Wild type and lcb1-100 cells were arrested in G1 at 30°C and released into media at restrictive temperature (37°C) and 10 *µ*M phytosphingosine was added 10 minutes after release from the arrest. Pkc1 phosphorylation and Clb2 protein levels were assayed by western blot. A background band from the same blot was used as a loading control.

The behavior of Pkc1 in the *lcb1-100* mutant presents a paradox: inhibition of Ypk1/2 caused a large loss of Pkc1 phosphorylation that was rescued by addition of phytosphingosine; however, the *lcb1-100* mutant caused only a partial loss of Pkc1 phosphorylation. There are several models that could explain this paradox. One is that the *lcb1-100* mutant causes only a partial inhibition of sphingolipid production. However, we think this is unlikely because we found that myriocin also caused only a partial loss of Pkc1 phosphorylation. Thus, another possibility is that inhibition of Ypk1/2 shuts down multiple pathways for production of phytosphingosine, whereas inactivation of Lcb1 only shuts down one pathway. For example, although phytosphingosine is synthesized by Lcb1, it can also be generated by dephosphorylation of phytosphingosine 1-phosphate (Dickson, 2008). Thus, when Lcb1 is inhibited, it is possible that the cell can temporarily compensate by generating phytosphingosine from other reserves, whereas inactivation of Ypk1/2 could shut down all pathways for production of phytosphingosine. It is also possible that inactivation of Ypk1/2 triggers pathways that lead to more rapid depletion of the sphingolipids that influence Pkc1 phosphorylation. Lastly, Lcb1 is in a complex with Sac1, which is a phosphatidylinositol-4-phosphate phosphatase (Breslow *et al*., 2010). Therefore, inactivation of Lcb1 could have effects on regulation of Sac1. This could also be true for inhibition of Lcb1 by myriocin because drug binding can affect protein structure in ways that have profound effects on protein function and protein-protein interactions (see for example, (Poulikakos *et al*., 2010)). Thus, inactivation of Lcb1 could lead to misregulation of Sac1, leading to effects on Pkc1. For example, normal regulation of Sac1 could be necessary for full activity of the phosphatase that dephosphorylates Pkc1, which would explain why Pkc1 is not fully dephosphorylated when Lcb1 is inhibited.

Together, the data indicate that an arrest of membrane growth causes an abrupt decrease in sphingolipid-dependent signals. The data further suggest that the TORC2-Ypk1/2 network monitors vesicle-dependent delivery of sphingolipids to the plasma membrane, rather than synthesis of sphingolipids at the endoplasmic reticulum. Finally, the data suggest that delivery of sphingolipids to the plasma membrane plays a role in controlling Pkc1 phosphorylation.

An interesting interpretation of the data is that the sphingolipid signaling network functions to coordinate aspects of cell growth. The network constitutes a feedback loop in which sphingolipids control TORC2 and Ypk1/2, while TORC2 and Ypk1/2 control production of sphingolipids. Sphingolipids are precursors for synthesis of components of the plasma membrane. Thus, the feedback loop could help ensure that growth of the plasma membrane is coordinated with synthesis of precursors at the endoplasmic reticulum. Since TORC2 and Ypk1/2 are also thought to control diverse aspects of cell growth, the feedback loop could work more broadly to coordinate events of cell growth. A previous study found that myriocin causes dose dependent hyperphosphorylation of the Orm1/2 proteins (Breslow et al., 2010). This observation suggests that the signaling network can measure the amount of sphingolipids transported to the plasma membrane, rather than simply measuring whether they are being transported. In this case, the network could generate signals that are proportional to the amount of sphingolipids transported to the plasma membrane. This kind of proportional growth signal could be used to measure and control cell size. Indeed, the data suggest that growth-dependent phosphorylation of Pkc1 may be dependent upon signals from sphingolipids.

The idea that cell growth and size are controlled by lipid-dependent signals is appealing. Control of cell growth and size would have been essential for survival of the earliest cells, so it is likely that the underlying mechanisms are ancient and conserved. Membranes that allow compartmentalization are one of the most fundamental and conserved features of cells, so it would make sense that mechanisms that control membrane growth evolved early and in close association with membrane lipids. Overall, we have a surprisingly limited understanding of the mechanisms and regulation of membrane growth during the cell cycle. A full understanding of cell growth, and the mechanisms that limit growth to control cell size, will require a deeper understanding of these mechanisms.

## Materials and Methods

### Yeast strains, culture conditions and plasmids

Most strains used in this study are in the W303 strain background (leu2-3,112 ura3-1 can1-100 ade2-1 his3-11,15 trp1-1 GAL+ ssd1-d2), except for DDY903 and YSY1269, which are in the S288C background (his3-Δ200 leu2-3, 112, ura3-52). The additional features of the strains used this study are listed in Table S5. Cells were grown in YEPD media (1% yeast extract, 2% peptone, 2% dextrose) supplemented with 40 mg/L adenine, or in YEP media (1% yeast extract, 2% peptone) supplemented with 40 mg/L adenine and with different carbon sources, as noted.

One-step PCR-based gene replacement was used for making deletions and adding epitope tags at the endogenous locus (oligonucleotide sequences available upon request) (Longtine et al., 1998; Janke et al., 2004). To make a strain that expresses an analog-sensitive allele of PKH1, the PKH1 gene was amplified and cloned into vector pRS306 (URA3). Site directed mutagenesis was then used to introduce mutations L203A and F187V to make plasmid pJZ5A (pkh1-as, URA3), as previously described (Sun et al., 2012). DK2547 was made by digesting plasmid pJZ5A with MscI to target integration at the PKH1 locus in *pkh2*Δ cells. After selection for plasmid integration, recombination events that loop out the plasmid were selected on FOA, and sequencing was used to identify looping out events that left the mutants behind. DK2876 was constructed similarly by digesting the vector pRS306-YPK1-as (ypk1-L424A URA3) with AflII, and integrating at YPK1 in *ypk2*Δ cells, followed by selection on URA3 and subsequent selection on FOA (Sun et al., 2012).

### Cell cycle time courses

To synchronize cells in G1 phase, cells were grown overnight to log phase in YEPD at room temperature. Cells at an OD600 of 0.6 were arrested in G1 by addition of alpha factor to 0.5 *µ*g/ml (bar1-strains) or 15 *µ*g/ml (BAR+ strains). Cells were arrested at room temperature for 3.5 hours and released by washing 3X with fresh YEPD. Time courses were carried out at 25°C unless otherwise noted. To prevent cells from re-entering the cell cycle, alpha factor was added back at 90 minutes after release from arrest. For time courses using analog-sensitive alleles, the cells were grown in YEPD media without supplemental adenine.

### Western blotting

To prepare samples for western blotting, 1.6 ml of culture were collected and centrifuged at 13000 rpm for 30s. The supernatant was removed and 250 *µ*l of glass beads were added before freezing in liquid nitrogen. Cells were lysed by bead-beating in 140*µ*l of 1X sample buffer (65 mM Tris-HCl, pH 6.8, 3% SDS, 10% glycerol, 50 mM NaF, 100 mM beta-glycerolphosphate, 5% 2-mercaptoethanol, 2 mM phenylmethylsulfonyl fluoride (PMSF) and bromophenol blue). The PMSF was added immediately before lysis from a 100 mM stock in ethanol. Cells were lysed in a mini-beadbeater-16 (Biospec Products) at top speed for 2 minutes. The samples were then centrifuged for 15 seconds at 13000 rpm, placed in boiling water bath for 5 min, and centrifuged for 5 min at 13000 rpm. SDS-PAGE was carried out as previously described using a 10% acrylamide gel (Harvey et al., 2005). Gels were run at a constant current of 20 mA until a prestained molecular weight marker of 57.6 kD was near the bottom of the gel.

For western blotting, protein was transferred to nitrocellulose membranes (or PVDF membranes for Pkc1 probing) for 1h 30min at 800 mA at 4°C in a Hoeffer transfer tank in buffer containing 20 mM Tris base, 150 mM glycine, and 20% methanol. Blots were probed overnight at room temperature with affinity-purified rabbit polyclonal antibodies raised against Pkc1, Clb2, or HA peptide (Anastasia et al., 2012). Anti-FLAG rabbit polyclonal antibody was from Sigma-Aldrich. TORC2-dependent phosphorylation of endogenous Ypk1 was detected using rabbit polyclonal anti-phospho-Ypk1(T662) (Gift of Ted Powers) and Pkh1/2-dependent phoshorylation of Ypk1 and Ypk2 was detected using a phospho-specific antibody that recognizes Ypk1 phosphorylated at T504 (Santa Cruz Biotechnology, Dallas, TX; Catalog number: sc-16744 P).

Phos-Tag gels were used to resolve phosphorylated forms of Orm2-Flag (Kinoshita et al., 2006). Samples were loaded onto 10% polyacrylamide gels containing 50 μM Phos-Tag (Wako Chemicals USA, Richmond, VA) and 100 μM MnCl2. Gels were rinsed twice for 10 min in transfer buffer containing 2 mM EDTA. Gels were transferred to nitrocellulose using the Trans-Blot Turbo^TM^ transfer system (BioRad USA, Hercules, CA).

### Analysis of bud emergence and polarity

To analyze bud emergence and polarity, samples from synchronized cultures were fixed with 3.7% formaldehyde and analyzed (n=200 cells) by light microscopy as previously described (Pringle et al., 1991). Bud polarity was analyzed in samples taken 120 minutes after release from G1 phase arrest. ImageJ was used to measure bud length and width, and the ratio was averaged for at least 50 cells in each sample. The average length and width of each strain were depicted as ovals (lines represent standard error) included with the bar graph whose relative shape and size can be directly compared.

### Preparation of samples for mass spectrometry

To prepare samples for mass spectrometry, wild type and sec6-4 cells were grown in YEPD medium overnight at room temperature to an optical density of O.D.600 0.8. Cells were arrested in G1 with mating pheromone and released from the arrest at room temperature. At 70 minutes after release from the G1 arrest, cells were shifted to a 34°C water bath and samples for mass spectrometry were taken 5 minutes after the shift. Samples for western blotting were collected before and after the shift to confirm that Pkc1 underwent dephosphorylation.

For each sample, 50 ml of cells were harvested by centrifuging 1 minute at 3800 rpm. The cells were resuspended in 1 ml YPD and transferred to a 2 ml screw-top tube. The cells were pelleted for 30 seconds, the supernatant was removed, and 250 ul of glass beads were added before freezing the cells on liquid nitrogen. All tubes, centrifuge buckets, and media were pre-warmed to 34°C to maintain restrictive temperature during cell collection.

To lyse the cells, 500 *µ*l of ice cold lysis buffer (8 M urea, 75 mM NaCl, 50 mM Tris-HCl pH 8.0, 50 mM NaF, 50 mM ß-glycerophosphate, 1 mM sodium orthovanadate, 10 mM sodium pyrophosphate, 1 mM PMSF) were added to the harvested cells before lysis using a Biospec Multibeater-16 at top speed for three cycles of 1 min, each followed by a 1 min incubation on ice to avoid over-heating. Samples were centrifuged at 13000 rpm for 10 minutes at 4°C and the lysates were transferred to fresh 1.6 ml tubes and spun again before transferring to new 1.6 ml tubes. A 5 *µ*L aliquot was taken from each sample for Bradford assay and protein quantification before flash freezing in liquid nitrogen. Three biological replicates for each strain were collected and analyzed by mass spectrometry.

Disulfide bonds were reduced by adding dithiothreitol to a final concentration of 2.5 mM and incubating at 56°C for 40 min. The extract was allowed to cool to room temperature and the reduced cysteines were alkylated by adding iodoacetamide to 7.5 mM and incubating for 40 min in the dark at room temperature. Alkylation was quenched with an additional 5 mM dithiothreitol. Peptide digestion and labeling by reductive dimethylation were carried out as previously described (Zapata et al., 2014).

### Phosphopeptide enrichment by SCX/TiO_2_

Phosphopeptides were enriched using a modified version of the two-step, SCX-IMAC/TiO_2_ protocol employing step elution from self-packed solid-phase extraction strong cation exchange (SCX) chromatography cartridges as previously described with some changes (Villén and Gygi, 2008). Peptides were resuspended in 1 ml SCX buffer A (7 mM KH_2_PO_4_, pH 2.65, 30% ACN) and loaded onto pre-equilibrated syringe-barrel columns packed with 500 mg of 20 *µ*m, 300 Å, polysulfoethylA resin (poly LC). The loading flow-through was collected and pooled with a 2 ml wash with buffer A. Seven additional fractions were collected after sequential addition of 3 ml of SCX buffer A containing increasing concentrations of KCl; 10, 20, 30, 40, 50, 60 and 100 mM KCl. All fractions were frozen in liquid nitrogen, lyophilized, resuspended in 1 ml of 1% FA, and desalted on 50 mg Sep-paks. Peptides were eluted with 500 *µ*l of 70% ACN, 1% FA. Five percent of each fraction was taken off for protein abundance analysis. The remaining peptides were dried in a speed vac. TiO_2_ enrichment was performed as in (Kettenbach and Gerber, 2011). Dried peptides were resuspended in 300 *µ*l wash/binding buffer (50% ACN, 2 M lactic acid) and incubated with 90 mg of prewashed Titansphere TiO_2_ beads (GL Sciences, #5020-75000) with vigorous shaking for 60 minutes at room temperature. The beads were washed two times with 300 *µ*l of wash/binding buffer and then two times with 300 ul 50% ACN/1% FA. Phosphopeptides were eluted in two steps by sequential treatments with 75 *µ*l 50 M KH_2_PO_4_, pH 10.7. The eluates were acidified by the addition of FA to 2% final concentration, desalted on STAGE tips (Rappsilber et al., 2003), and dried in a speed vac. Eight fractions were analyzed by LC-MS/MS.

### Mass spectrometry

Phosphopeptide samples were analyzed on a LTQ Orbitrap Velos mass spectrometer (Thermo Fisher Scientific) equipped with an Accela 600 quaternary pump (Thermo Fisher Scientific) and a Famos microautosampler (LC Packings, Sunnyvale, CA). Nanospray tips were hand-pulled using 100 μm I.D. fused-silica tubing and packed with 0.5 cm of Magic C4 resin (5μm, 100 Å, Michrom Bioresources, Auburn, CA) followed by 20 cm of Maccel C18AQ resin (3μm, 200 Å, Nest Group, Southborough, MA). Peptides were separated using a gradient of 3% to 28% ACN in 0.125% FA over 70 min with an on column flow rate of ∼300-500 nl/min.

Peptides were detected using a data-dependent Top20-MS2 method. For each cycle, one full MS scan of m/z = 300-1500 was acquired in the Orbitrap at a resolution of 60,000 at m/z = 400 with AGC target = 1 × 10^6^ and maximum ion accumulation time of 500 mS. Each full scan was followed by the selection of the most intense ions, up to 20, for collision induced dissociation (CID) and MS2 analysis in the LTQ. An AGC target of 2 × 10^3^ and maximum ion accumulation time of 150 mS was used for MS2 scans. Ions selected for MS2 analysis were excluded from re-analysis for 60 sec. Precursor ions with charge = 1+ or unassigned were excluded from selection for MS2 analysis. Lockmass, employing atmospheric polydimethylsiloxane (m/z = 445.120025) as an internal standard was used in all runs to calibrate orbitrap MS precursor masses. Each sample was analyzed twice for a total of 48 runs.

### Peptide identification and filtering

MS2 spectra were searched using SEQUEST v.28 (rev. 13) (Eng et al., 1994) against a composite database containing the translated sequences of all predicted open reading frames of Saccharomyces cerevisiae (http://downloads.yeastgenome.org, downloaded 10/30/2009) and its reversed complement, using the following parameters: a precursor mass tolerance of ±20 ppm; 1.0 Da product ion mass tolerance; lysC digestion; up to two missed cleavages; static modifications of carbamidomethylation on cysteine (+57.0214), dimethyl adducts (+28.0313) on lysine and peptide amino termini; and dynamic modifications for methionine oxidation (+15.9949), heavy dimethylation (+6.0377) on lysine and peptide amino termini, and phosphate (+79.9663) on serine, threonine, and tyrosine for phosphopeptide enriched samples.

Peptide spectral matches were filtered to 1% false discovery rate (FDR) using the target-decoy strategy (Elias and Gygi, 2007) combined with linear discriminant analysis (LDA) (Huttlin et al., 2010) using several different parameters including Xcorr, ΔCn’, precursor mass error, observed ion charge state, and predicted solution charge state. Linear discriminant models were calculated for each LC-MS/MS run using peptide matches to forward and reversed protein sequences as positive and negative training data. Peptide spectral matches within each run were sorted in descending order by discriminant score and filtered to a 1% FDR as revealed by the number of decoy sequences remaining in the data set. The data were further filtered to control protein level FDRs. Peptides from all fractions in each experiment were combined and assembled into proteins. Protein scores were derived from the product of all LDA peptide probabilities, sorted by rank, and filtered to 1% FDR as described for peptides. Protein filtering removes many additional decoy peptides hits, thus further reducing the peptide FDR. The final peptide FDR was 0.08%. The remaining 65 peptide matches to the decoy database were removed from the final dataset.

Mass spectrometric feature detection and peptide quantification were performed with custom software developed in the Gygi Lab (Bakalarski *et al*., 2007). Peptide peak noise levels were calculated as the median peak intensity of all peaks +/−10 Th surrounding the precursor m/z in a window of +/−20 MS1 scans. Signal-to-noise was calculated as the ratio of the instrument reported precursor m/z peak intensity to the calculated noise value. For inclusion in quantitative calculations, peptides were required to have a sum signal-to-noise ratio ≥ 10 for heavy and light species. Ratios were normalized to recenter the distribution at 1:1 (log_2_ = 0). Due to the sparsity of the data and the frequent uncertainty of site localization that is inherent in these analyses, we grouped same sequence, same phosphorylation site number peptides together for quantification. Thus, phosphorylation site ratios were calculated for each replicate from the median of all such peptides.

Phosphorylation site localization analysis was done using the Ascore algorithm (Beausoleil et al., 2006). These values appear in Tables S1 and S2. Sites with Ascore ≥ 13 were considered localized. Because localization and quantification are only weakly correlated, for the purposes of quantification, unlocalized site assignments were reassigned using assignments corresponding to the maximum Ascore from other same sequence peptides in the dataset.

All raw data files, peak lists, and the sequence database have been deposited in the MASSive repository and are available for download at ftp://massive.ucsd.edu/MSV000080919.

## Acknowledgements

We thank Ted Powers for anti-phospho-Ypk1(T662P) antibody and François Roelants for advice and critical reading of the manuscript. We also thank Yidi Sun and David Drubin for advice and sharing strains, members of the Kellogg laboratory for advice and critical reading of the manuscript, and Ben Abrams for assistance with microscopy. This work was supported by NIH Grant GM109143.

## Competing Interests

The authors have no competing interests

## List of Tables

**Table S1**

All identified phosphorylation sites.

**Table S2**

All identified phosphopeptides.

**Table S3**

Sites that show a significant decrease in phosphorylation in response to an arrest of membrane growth.

**Table S4**

Sites that show a significant increase in phosphorylation in response to an arrest of membrane growth.

**Table S5**

Yeast strains used in this study.

**Figure Supplement 1.**
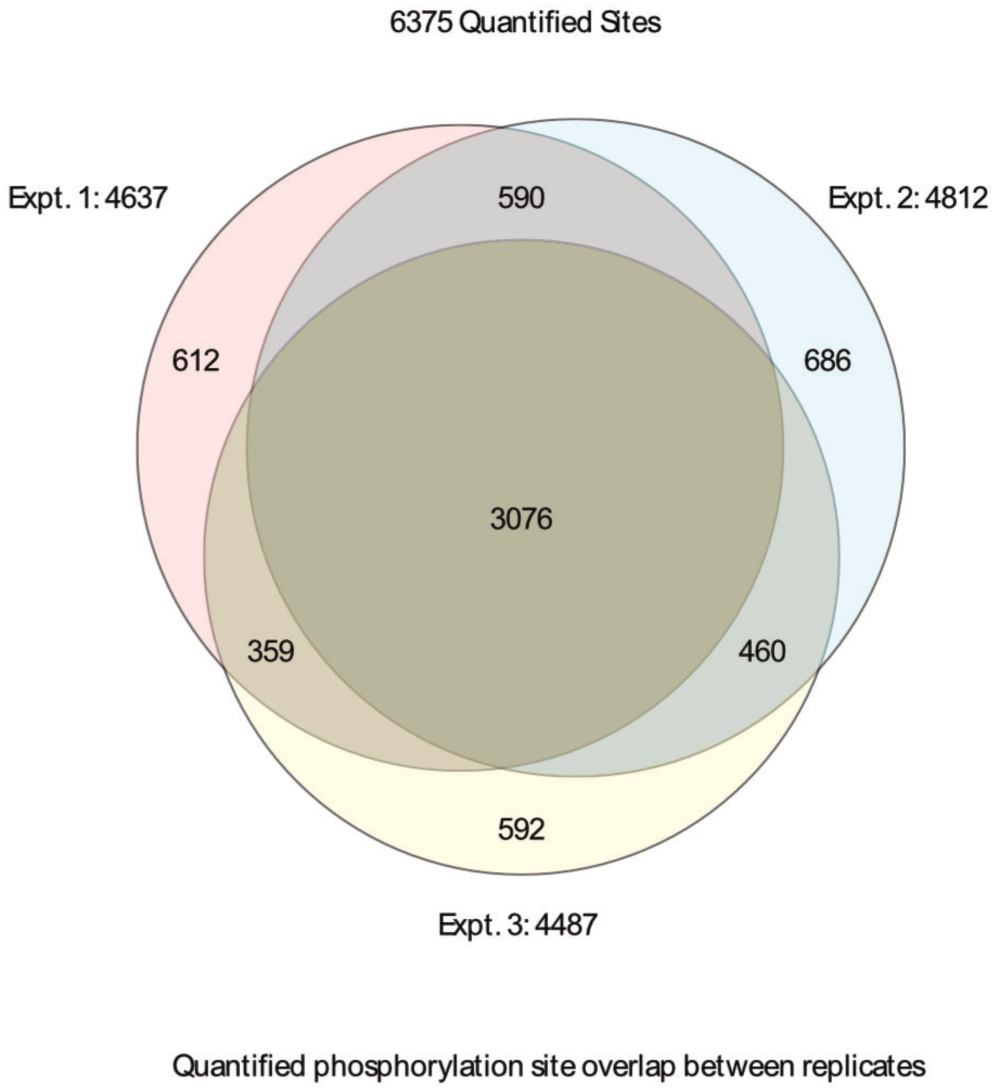
Overlap of quantified sites across three biological replicates. All quantified sites and their distribution and overlap are depicted.

**Figure Supplement 2.**
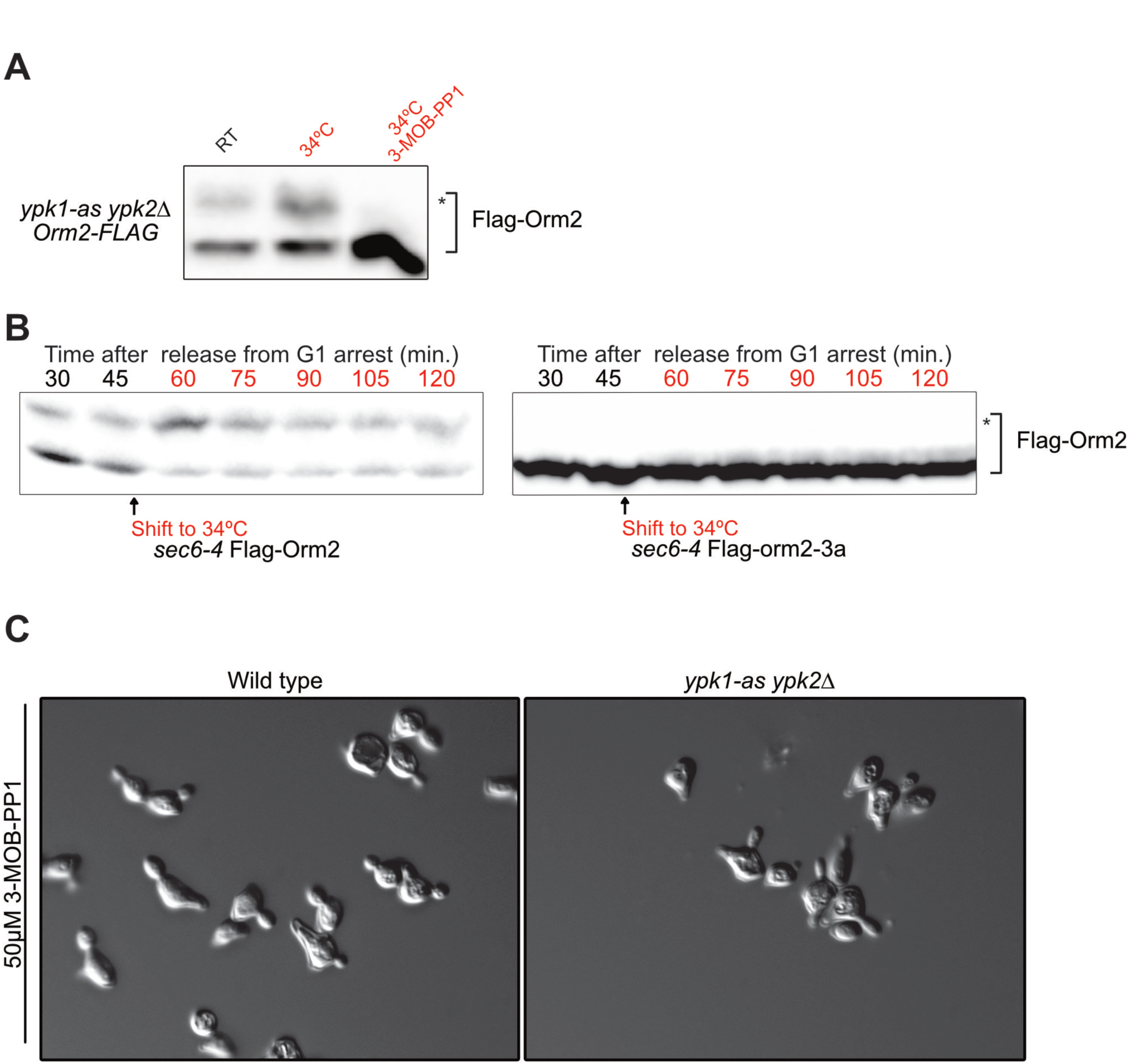
(**A**) ypk1-as *ypk2*Δ sec6-4 Flag-ORM2 cells were released from a G1 arrest and split into three identical cultures. At 59 minutes after release, 3-MOB-PP1 was added to one culture and vehicle (DMSO) was added to the others. At 60 minutes, the culture containing 3-MOB-PP1 was shifted to 34C along with one of the other cultures. A control culture was left at room temperature. Samples were taken from all three cultures at 75 minutes and Orm2 phosphorylation was detected with a phos-tag gel. In this experiment, hyperphosphorylation of Flag-Orm2 in response to inactivation of Sec6 was diminished compared to Figure 1E, which may be due to reduced activity of the ypk1-as allele. **(B)** sec6-4 Flag-ORM2 and sec6-4 Flag-orm2-3a (S46A, S47A, S48A) cells were released from a G1 arrest. After the 45 minute time point both strains were shifted to 34°C. Orm2 phosphorylation was detected with a phos-tag gel and western blotting. **(C)** Wild type and ypk1-as *ypk2*Δ cells were released from a G1 arrest and 50 *µ*M 3-MOB-PP1 inhibitor was added at 10 minutes. Cells at the 120 minute time point were fixed and imaged by DIC microscopy.

**Figure Supplement 3.**
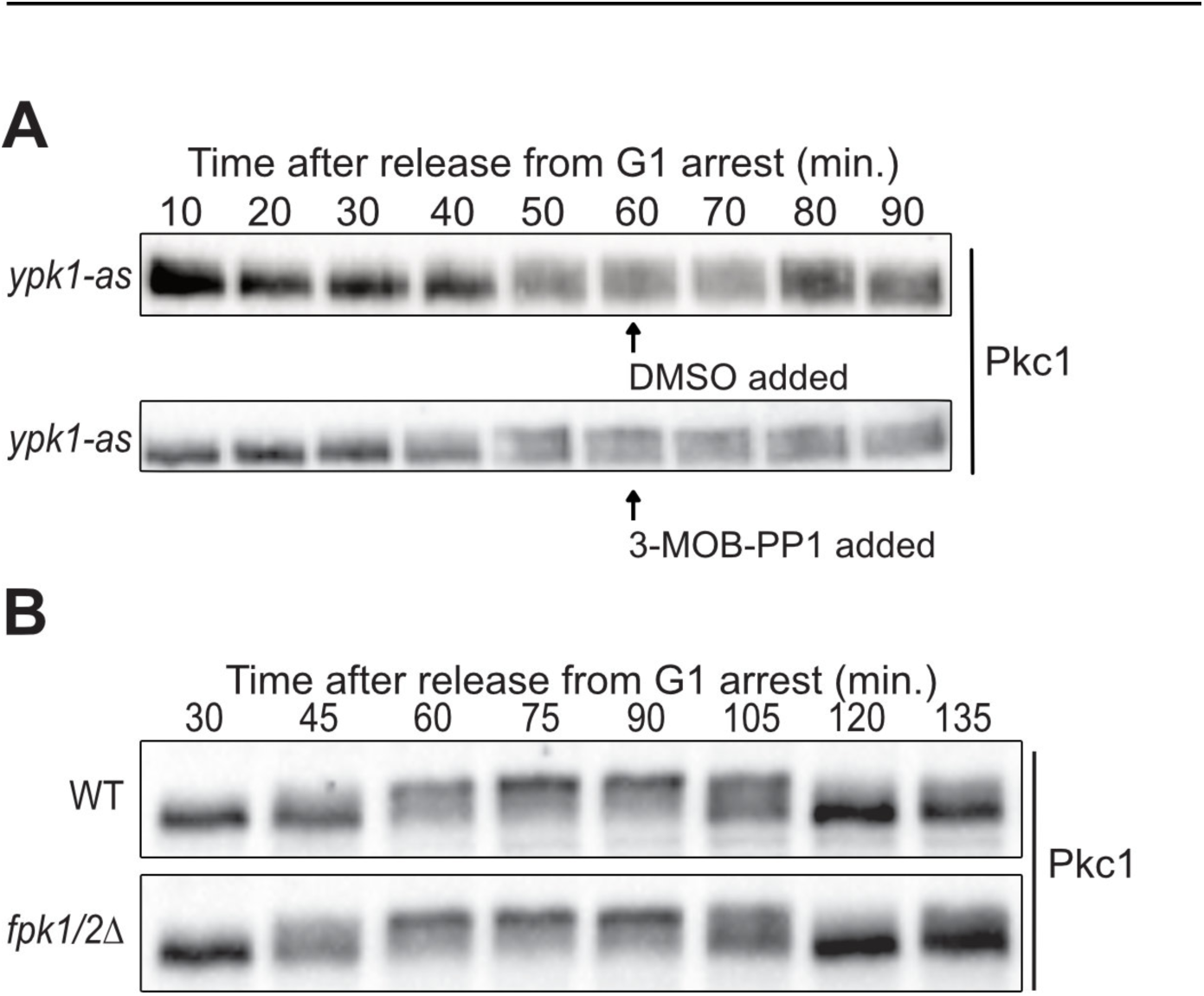
(**A**) ypk1-as *ypk2*Δ cells were released from a G1 arrest at 25°C and the indicated compounds were added at 60 minutes after release when small buds were present. Pkc1 phosphorylation was assayed by western blot. (**B**) Wild type and *fpk1/2*Δ cells were released from a G1 arrest at 25°C. Pkc1 phosphorylation was assayed by western blot.

## References

Abe, M., Qadota, H., Hirata, A., and Ohya, Y. (2003). Lack of GTP-bound Rho1p in secretory vesicles of Saccharomyces cerevisiae. The Journal of Cell Biology 162, 85–97.

Anastasia, S. D., Nguyen, D. L., Thai, V., Meloy, M., MacDonough, T., and Kellogg, D. R. (2012). A link between mitotic entry and membrane growth suggests a novel model for cell size control. The Journal of Cell Biology 197, 89–104.

Andrews, P. D., and Stark, M. J. (2000). Dynamic, Rho1p-dependent localization of Pkc1p to sites of polarized growth. Journal of Cell Science 113 (Pt 15), 2685–2693.

Aronova, S., Wedaman, K., Aronov, P. A., Fontes, K., Ramos, K., Hammock, B. D., and Powers, T. (2008). Regulation of Ceramide Biosynthesis by TOR Complex 2. Cell Metabolism 7, 148–158.

Bakalarski, C. E., Haas, W., Dephoure, N. E., and Gygi, S. P. (2007). The effects of mass accuracy, data acquisition speed, and search algorithm choice on peptide identification rates in phosphoproteomics. Anal Bioanal Chem 389, 1409–1419.

Berchtold, D., and Walther, T. C. (2009). TORC2 plasma membrane localization is essential for cell viability and restricted to a distinct domain. Mol. Biol. Cell 20, 1565–1575.

Berchtold, D., Piccolis, M., Chiaruttini, N., Riezman, I., Riezman, H., Roux, A., Walther, T. C., and Loewith, R. (2012). Plasma membrane stress induces relocalization of Slm proteins and activation of TORC2 to promote sphingolipid synthesis. Nature Publishing Group 14, 542–547.

Breslow, D. K., Collins, S. R., Bodenmiller, B., Aebersold, R., Simons, K., Shevchenko, A., Ejsing, C. S., and Weissman, J. S. (2010). Orm family proteins mediate sphingolipid homeostasis. Nature 463, 1048–1053.

Casamayor, A., Torrance, P. D., Kobayashi, T., Thorner, J., and Alessi, D. R. (1999). Functional counterparts of mammalian protein kinases PDK1 and SGK in budding yeast. Curr. Biol. 9, 186– 197.

Dickson, R. C. (2008). Thematic review series: sphingolipids. New insights into sphingolipid metabolism and function in budding yeast. The Journal of Lipid Research 49, 909–921.

Ejsing, C. S., Sampaio, J. L., Surendranath, V., Duchoslav, E., Ekroos, K., Klemm, R. W., Simons, K., and Shevchenko, A. (2009). Global analysis of the yeast lipidome by quantitative shotgun mass spectrometry. Proc. Natl. Acad. Sci. U.S.a. 106, 2136–2141.

Enciso, G., Kellogg, D. R., and Vargas, A. (2014). Compact modeling of allosteric multisite proteins: application to a cell size checkpoint. PLoS Comput Biol 10, e1003443.

Friant, S., Lombardi, R., Schmelzle, T., Hall, M. N., and Riezman, H. (2001). Sphingoid base signaling via Pkh kinases is required for endocytosis in yeast. Embo J 20, 6783–6792.

Friant, S., Zanolari, B., and Riezman, H. (2000). Increased protein kinase or decreased PP2A activity bypasses sphingoid base requirement in endocytosis. Embo J 19, 2834–2844.

Hatakeyama, R., Kono, K., and Yoshida, S. (2017). Ypk1 and Ypk2 kinases maintain Rho1 at the plasma membrane by flippase-dependent lipid remodeling after membrane stresses. Journal of Cell Science 130, 1169–1178.

Helliwell, S. B., Schmidt, A., Ohya, Y., and Hall, M. N. (1998). The Rho1 effector Pkc1, but not Bni1, mediates signalling from Tor2 to the actin cytoskeleton. Current Biology 8, 1211–1214.

Hua, Z., Fatheddin, P., and Graham, T. R. (2002). An essential subfamily of Drs2p-related P-type ATPases is required for protein trafficking between Golgi complex and endosomal/vacuolar system. Mol. Biol. Cell 13, 3162–3177.

Inagaki, M., Schmelzle, T., Yamaguchi, K., Irie, K., Hall, M. N., and Matsumoto, K. (1999). PDK1 homologs activate the Pkc1-mitogen-activated protein kinase pathway in yeast. Molecular and Cellular Biology 19, 8344–8352.

Janke, C. et al. (2004). A versatile toolbox for PCR-based tagging of yeast genes: new fluorescent proteins, more markers and promoter substitution cassettes. Yeast 21, 947–962.

Kamada, Y., Fujioka, Y., Suzuki, N. N., Inagaki, F., Wullschleger, S., Loewith, R., Hall, M. N., and Ohsumi, Y. (2005). Tor2 Directly Phosphorylates the AGC Kinase Ypk2 To Regulate Actin Polarization. Molecular and Cellular Biology 25, 7239–7248.

Kamada, Y., Qadota, H., Python, C. P., Anraku, Y., Ohya, Y., and Levin, D. E. (1996). Activation of yeast protein kinase C by Rho1 GTPase. J. Biol. Chem. 271, 9193–9196.

Kettenbach, A. N., and Gerber, S. A. (2011). Rapid and reproducible single-stage phosphopeptide enrichment of complex peptide mixtures: application to general and phosphotyrosine-specific phosphoproteomics experiments. Anal. Chem. 83, 7635–7644.

Klose, C., Surma, M. A., Gerl, M. J., Meyenhofer, F., Shevchenko, A., and Simons, K. (2012). Flexibility of a eukaryotic lipidome--insights from yeast lipidomics. PLoS ONE 7, e35063.

Lew, D. J., and Reed, S. I. (1993). Morphogenesis in the yeast cell cycle: regulation by Cdc28 and cyclins. The Journal of Cell Biology 120, 1305–1320.

Li, Y., Moir, R. D., Sethy-Coraci, I. K., Warner, J. R., and Willis, I. M. (2000). Repression of Ribosome and tRNA Synthesis in Secretion-Defective Cells Is Signaled by a Novel Branch of the Cell Integrity Pathway. Molecular and Cellular Biology 20, 3843–3851.

Liu, K., Zhang, X., Lester, R. L., and Dickson, R. C. (2005). The Sphingoid Long Chain Base Phytosphingosine Activates AGC-type Protein Kinases in Saccharomyces cerevisiae Including Ypk1, Ypk2, and Sch9. Journal of Biological Chemistry 280, 22679–22687.

Longtine, M. S., McKenzie, A., DeMarini, D. J., Shah, N. G., Wach, A., Brachat, A., Philippsen, P., and Pringle, J. R. (1998). Additional modules for versatile and economical PCR-based gene deletion and modification in Saccharomyces cerevisiae. Yeast 14, 953–961.

McCusker, D., and Kellogg, D. R. (2012). Plasma membrane growth during the cell cycle: unsolved mysteries and recent progress. Current Opinion in Cell Biology 24, 845–851.

Meier, K. D., Deloche, O., Kajiwara, K., Funato, K., and Riezman, H. (2006). Sphingoid base is required for translation initiation during heat stress in Saccharomyces cerevisiae. Mol. Biol. Cell 17, 1164–1175.

Mizuta, K., and Warner, J. R. (1994). Continued functioning of the secretory pathway is essential for ribosome synthesis. Molecular and Cellular Biology 14, 2493–2502.

Muir, A., Ramachandran, S., Roelants, F. M., Timmons, G., and Thorner, J. (2014). TORC2-dependent protein kinase Ypk1 phosphorylates ceramide synthase to stimulate synthesis of complex sphingolipids. eLife 3.

Nakano, K., Yamamoto, T., Kishimoto, T., Noji, T., and Tanaka, K. (2008). Protein kinases Fpk1p and Fpk2p are novel regulators of phospholipid asymmetry. Mol. Biol. Cell 19, 1783– 1797.

Nanduri, J., and Tartakoff, A. M. (2001). The arrest of secretion response in yeast: signaling from the secretory path to the nucleus via Wsc proteins and Pkc1p. Molecular Cell 8, 281–289.

Niles, B. J., and Powers, T. (2014). TOR complex 2-Ypk1 signaling regulates actin polarization via reactive oxygen species. Mol. Biol. Cell 25, 3962–3972.

Niles, B. J., Mogri, H., Hill, A., Vlahakis, A., and Powers, T. (2012). Plasma membrane recruitment and activation of the AGC kinase Ypk1 is mediated by target of rapamycin complex 2 (TORC2) and its effector proteins Slm1 and Slm2. Proc. Natl. Acad. Sci. U.S.a. 109, 1536– 1541.

Nonaka, H., Tanaka, K., Hirano, H., Fujiwara, T., Kohno, H., Umikawa, M., Mino, A., and Takai, Y. (1995). A downstream target of RHO1 small GTP-binding protein is PKC1, a homolog of protein kinase C, which leads to activation of the MAP kinase cascade in Saccharomyces cerevisiae. Embo J 14, 5931–5938.

Olson, D. K., Fröhlich, F., Christiano, R., Hannibal-Bach, H. K., Ejsing, C. S., and Walther, T. C. (2015). Rom2-dependent phosphorylation of Elo2 controls the abundance of very long-chain fatty acids. J. Biol. Chem. 290, 4238–4247.

Pichler, H., Gaigg, B., Hrastnik, C., Achleitner, G., Kohlwein, S. D., Zellnig, G., Perktold, A., and Daum, G. (2001). A subfraction of the yeast endoplasmic reticulum associates with the plasma membrane and has a high capacity to synthesize lipids. Eur. J. Biochem. 268, 2351–2361.

Pomorski, T., Lombardi, R., Riezman, H., Devaux, P. F., van Meer, G., and Holthuis, J. C. M. (2003). Drs2p-related P-type ATPases Dnf1p and Dnf2p are required for phospholipid translocation across the yeast plasma membrane and serve a role in endocytosis. Mol. Biol. Cell 14, 1240–1254.

Poulikakos, P. I., Zhang, C., Bollag, G., Shokat, K. M., and Rosen, N. (2010). nature08902. Nature 464, 427–430.

Roelants, F. M., Baltz, A. G., Trott, A. E., Fereres, S., and Thorner, J. (2010). A protein kinase network regulates the function of aminophospholipid flippases. Proc. Natl. Acad. Sci. U.S.a. 107, 34–39.

Roelants, F. M., Breslow, D. K., Muir, A., Weissman, J. S., and Thorner, J. (2011). Protein kinase Ypk1 phosphorylates regulatory proteins Orm1 and Orm2 to control sphingolipid homeostasis in Saccharomyces cerevisiae. Proc. Natl. Acad. Sci. U.S.a. 108, 19222–19227.

Roelants, F. M., Torrance, P. D., and Thorner, J. (2004). Differential roles of PDK1- and PDK2- phosphorylation sites in the yeast AGC kinases Ypk1, Pkc1 and Sch9. Microbiology (Reading, Engl.) 150, 3289–3304.

Roelants, F. M., Torrance, P. D., Bezman, N., and Thorner, J. (2002). Pkh1 and Pkh2 differentially phosphorylate and activate Ypk1 and Ykr2 and define protein kinase modules required for maintenance of cell wall integrity. Mol. Biol. Cell 13, 3005–3028.

Rossio, V., and Yoshida, S. (2011). Spatial regulation of Cdc55-PP2A by Zds1/Zds2 controls mitotic entry and mitotic exit in budding yeast. The Journal of Cell Biology 193, 445–454.

Schmelzle, T., Helliwell, S. B., and Hall, M. N. (2002). Yeast Protein Kinases and the RHO1 Exchange Factor TUS1 Are Novel Components of the Cell Integrity Pathway in Yeast. Molecular and Cellular Biology 22, 1329–1339.

Sun, Y., Miao, Y., Yamane, Y., Zhang, C., Shokat, K. M., Takematsu, H., Kozutsumi, Y., and Drubin, D. G. (2012). Orm protein phosphoregulation mediates transient sphingolipid biosynthesis response to heat stress via the Pkh-Ypk and Cdc55-PP2A pathways. Mol. Biol. Cell 23, 2388–2398.

Sun, Y., Taniguchi, R., Tanoue, D., Yamaji, T., Takematsu, H., Mori, K., Fujita, T., Kawasaki, T., and Kozutsumi, Y. (2000). Sli2 (Ypk1), a homologue of mammalian protein kinase SGK, is a downstream kinase in the sphingolipid-mediated signaling pathway of yeast. Molecular and Cellular Biology 20, 4411–4419.

Villén, J., and Gygi, S. P. (2008). The SCX/IMAC enrichment approach for global phosphorylation analysis by mass spectrometry. Nat Protoc 3, 1630–1638.

Wullschleger, S., Loewith, R., Oppliger, W., and Hall, M. N. (2005). Molecular organization of target of rapamycin complex 2. J. Biol. Chem. 280, 30697–30704.

Yamochi, W., Tanaka, K., Nonaka, H., Maeda, A., Musha, T., and Takai, Y. (1994). Growth site localization of Rho1 small GTP-binding protein and its involvement in bud formation in Saccharomyces cerevisiae. The Journal of Cell Biology 125, 1077–1093.

